# Generation time and seasonal migration explain variation in spatial population synchrony across European bird species

**DOI:** 10.1101/2023.01.30.526164

**Authors:** Ellen C. Martin, Brage Bremset Hansen, Aline Magdalena Lee, Ivar Herfindal

## Abstract

1. Spatial population synchrony is common among populations of the same species and is an important predictor of extinction risk. Despite the potential consequences for metapopulation persistence, we still largely lack understanding of what makes one species more likely to be synchronized than another given the same environmental conditions.
2. Generally, environmental conditions on a shared environment or a species’ sensitivity to the environment can explain the extent of synchrony. Populations that are closer together experience more similar fluctuations in their environments than those populations that are further apart and are therefore more synchronized. The relative importance of environmental and demographic stochasticity for population dynamics is strongly linked to species’ life history traits, such as pace of life, why may impact population synchrony. For populations that migrate, there may be multiple environmental conditions at different locations driving synchrony. However, the importance of life history and migration strategies in determining patterns of spatial population synchrony have rarely been explored empirically. We therefore hypothesize that generation time, a proxy for pace of life, and migration play an important role in determining spatial population synchrony.
3. We used population abundance data on breeding birds from four countries to investigate patterns of spatial population synchrony in growth rate and abundance. We investigated differences in synchrony across a gradient of generation times in resident, short-distance migrant, and long-distance migrant bird species.
4. Species with shorter generation times were more synchronized than species with longer generation times. Short-distance migrants were more synchronized than long-distance migrants and resident birds.
5. Our results provide novel empirical links between spatial population synchrony and species traits known to be of key importance for population dynamics, generation time and migration characteristics. We show how these different mechanisms can be combined to understand species-specific causes of spatial population synchrony. Understanding these specific drivers of spatial population synchrony is important in the face of increasingly severe threats to biodiversity and could be key for successful future conservation outcomes.

## Introduction

Spatial population synchrony, i.e., the correlated fluctuation of population abundances in different places, is common between populations of the same species and an important predictor of extinction risk, since metapopulations composed of synchronized populations are more likely to go extinct (Heino et al., 1997). Synchrony has been identified in a wide number of taxa including insects, fish, birds, and mammals (e.g., Hanski et al., 1995, Ranta et al., 1995, Ims & Andreassen, 2000, Raimondo et al., 2004, Koenig, 2006, Jones et al., 2007, Sæther et al., 2007, Chevalier et al., 2014, Koenig & Liebhold, 2016, Hansen et al., 2019, Marquez et al., 2019). Despite the potential consequences for species persistence and the importance for conservation, we still largely lack understanding of which traits make one species more likely to be synchronized than another. We hypothesize that traits that determine the environments individuals are exposed to and traits that influence their sensitivities to those environments play an important role in determining their spatial population synchrony.

Spatial population synchrony has three main drivers: Correlated fluctuations in the environment (i.e., the Moran effect; Moran, 1953), individual movement (i.e., dispersal) between populations (Lande et al., 1999, Paradis et al., 1999), and interactions of individuals through spatially linked populations, such as a shared predator (Myrberget, 1973, Ims & Andreassen, 2000). These three drivers can impact both the scaling (i.e., the relationship between synchrony and distance) and mean spatial population synchrony (Kendall et al., 2000, Engen & Sæther, 2005). Stochastic variability over time and space in population dynamics is caused by environmental stochasticity, acting on all individuals similarly, and demographic stochasticity, i.e. random variation in survival and reproduction among individuals (Lande et al., 2003). Nearby populations experience more similar fluctuations (i.e., stochasticity) in the environment, and therefore higher population synchrony, than those populations which are further apart (Sæther, 1997, Lande et al., 1999, Ellis & Schneider, 2008). Species whose dynamics are more sensitive to environmental stochasticity would be expected to be more synchronized because they tend to have more immediate responses to environmental stochasticity. Unlike environmental stochasticity, demographic stochasticity is not autocorrelated in space, resulting in a decoupling of species’ dynamics from the environment in the presence of high demographic stochasticity (Engen & Sæther, 2016). The relative importance of environmental and demographic stochasticity for population dynamics is strongly linked to species’ life history traits (Lande et al., 2002, Sæther et al., 2013), and understanding the relationship between species traits and synchrony can help to understand differences in synchrony among species.

Life history can be considered as a gradient along which species are placed according to a trade-off in their traits, often called the slow-fast life history continuum, with high reproduction on one end and high survival on the other (Stearns, 1999). Generation time is often used as a proxy for multiple correlated traits along this slow-fast life history continuum, such as age at first reproduction, fecundity, and survival (Gaillard et al., 2005), and has successfully been used to describe patterns in population fluctuations (Marquez et al., 2019). Species with short generation times typically have high reproductive rates, low survival, and are on the fast end of the slow-fast life history continuum (i.e., r-selected species; MacArthur & Wilson, 1967), whereas species with longer generation times typically have low reproductive rates, higher survival, and are on the slow end of the slow-fast life history continuum (i.e., K-selected species; MacArthur & Wilson, 1967, Sæther & Bakke, 2000). Theoretical and empirical examples show that species with different generation times have different sensitivities to environmental variation (Tedesco & Hugueny, 2006, Bjørkvoll et al., 2012, Sæther et al., 2013, Chevalier et al., 2014), and that environmental stochasticity has a greater effect on population dynamics for species with shorter generation times (Sæther et al., 2005, Sæther et al., 2013). Some studies found evidence that generation time was related to the scaling of spatial population synchrony, where species with longer generation time had more synchronized dynamics over greater distances than those of species with shorter generation time (Marquez et al., 2019). Furthermore, species with different generation times have different sensitivities in their abundances and population growth rates to demographic stochasticity (Sæther et al., 2013, Marquez et al., 2019). Species with longer generation times typically have smaller population abundances, which can result in a larger effect of demographic stochasticity on their dynamics (Sæther & Bakke, 2000, Ferguson & Larivière, 2002, Oli, 2004). Investigating if there is a relationship between contrasting life histories – and associated sensitivities to demographic and environmental stochasticity – with variation in spatial population synchrony is an important next step in understanding drivers and implications of such synchrony.

Space use and movement are important drivers of spatial population synchrony. Because individuals tend to move, the environment experienced varies not only because of temporal environmental stochasticity. Most studies on individual movement effects have focused on dispersal, finding that frequent dispersal, defined as a one-way movement which links population dynamics in spatially separate populations (e.g., Engen et al., 2002), synchronizes populations (Sutcliffe et al., 1996, Swanson & Johnson, 1999). However, two-way movement such as seasonal migration between different locations is a common phenomenon in nature that complicates studies of population dynamics but has huge implications for biodiversity and ecosystem functioning (Bauer & Hoye, 2014). Seasonal migration, the regular and reversible movement between locations across seasons typically between a nonbreeding ground and breeding ground (Webster et al., 2002, Somveille et al., 2021), often goes overlooked when considering drivers of spatial population synchrony. Migrating populations are exposed to several different environments through migratory routes and nonbreeding grounds (Newton, 2008), and these different environment and climate patterns are known to impact vital rates (Bogdanova et al., 2011, Selonen et al., 2021), either immediately or in the future as carry-over effects (Harrison et al., 2010). Species’ life history and sensitivity to environmental and demographic stochasticity may modify the consequences of such variation in migratory strategy on synchrony by rendering some species more sensitive to the different environments experienced through migration.

In this study, we explored the implications of two key life history traits – generation time and migration tactic – for spatial population synchrony across 94 bird species from four countries in Europe. Given known differences in sensitivities to environmental and demographic stochasticity among species with different life history traits, we expected higher synchrony for species with fast versus slow life histories, i.e., short versus long generation times, due to higher and lower sensitivities to environmental and demographic stochasticity respectively. We also expected that species that spent less time in correlated environments on the breeding ground, traveled further, and were exposed to more environmental stochasticity because individuals are more spread out in space (i.e., long-distance migrants) would be less synchronized than species that spent more time in one constant environment (i.e., resident species). We expected to see a gradient in increasing synchrony from long-distance migrants to short-distance migrants and resident species.

## Materials and Methods

### Study Area and data

We used population abundance data of breeding birds from four long-term monitoring programs in Norway, Sweden, Switzerland, and the United Kingdom. All surveys were conducted during the breeding season, between spring and mid-summer (Fig. 1).

**Figure 1.**
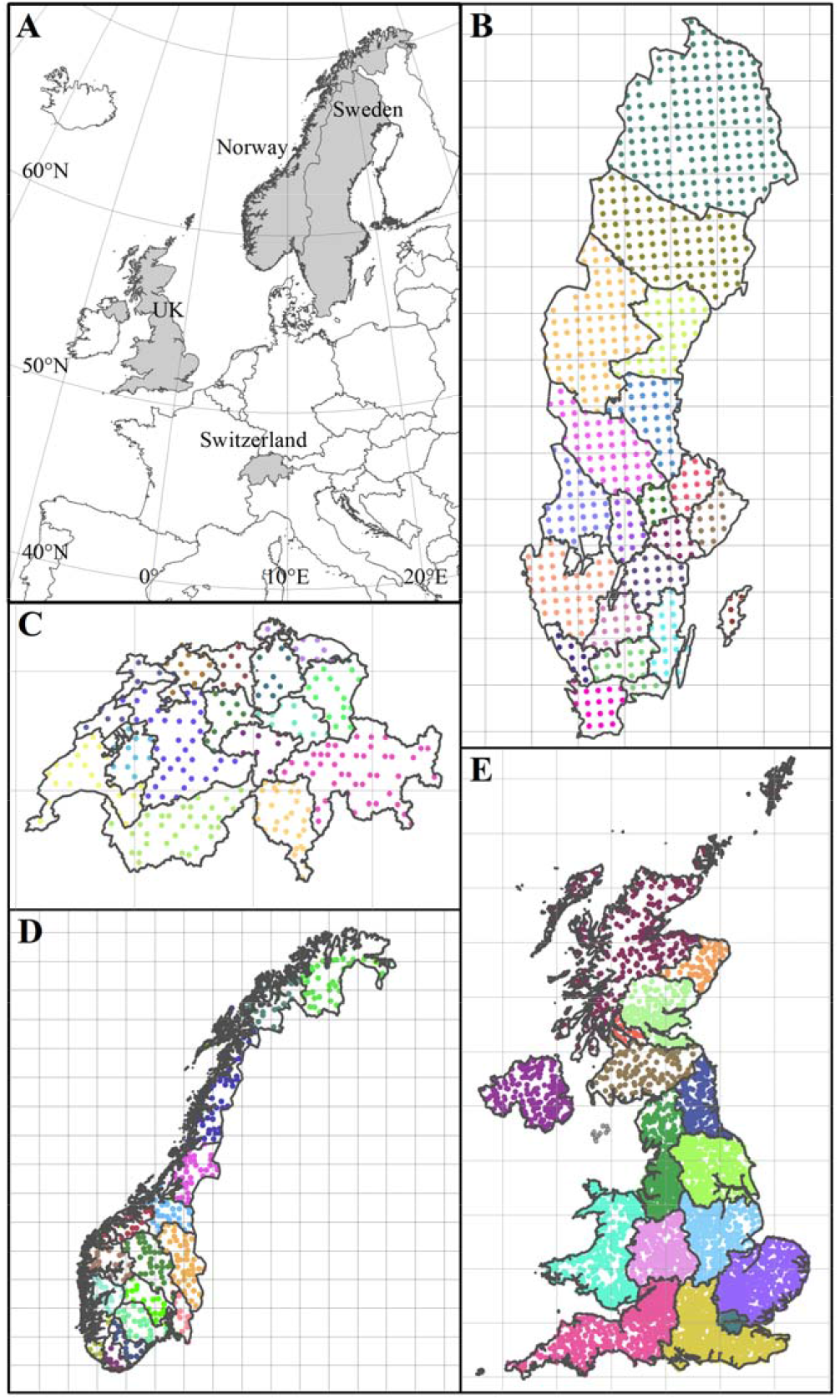
(A) The four study countries. Survey locations in each country presented in B-E. Administrative units were clustered with next nearest neighbor with fewest survey points to achieve a minimum of 8 sample locations. Black boundaries represent aggregated administrative unit boundaries. The grids in the country maps are 100 x100 km. Dots are survey locations, and the dot color represents which survey points are aggregated within each administrative unit. (B) Sweden, (C) Switzerland, (D) Norway, and (E) the United Kingdom.

#### Norway

Data were downloaded in September 2021 from the Global Biodiversity Information Facility (GBIF) with supplemental location and survey information provided by the Norwegian Institute for Nature Research (Kålås et al., 2022). Data were collected as a part of the Norwegian TOV-E Bird Survey and spanned years 2006 to 2021. The survey methodology involved conducting five-minute point count surveys within a 2km by 2km square (Kålås et al., 2022). Observers recorded all pairs of birds seen or heard during the transects. 494 unique survey points were surveyed over 19 years (Fig. 1D).

#### Sweden

Data were downloaded in March 2021 from GBIF (Lindström & Green, 2021). Data were from the *Swedish Bird Survey standardrutterna* (i.e., standardized fixed routes) line survey transects published by the Department of Biology at Lund University, and spanned years 2006 to 2019. The survey methodology involved conducting a fixed route survey of eight 1km-line transects within a 2km by 2km square (Lindström & Green, 2021). Observers recorded all birds seen or heard during the transects. 716 unique points were surveyed (Fig. 1B).

#### Switzerland

Data were provided in September 2020 by the Swiss Ornithological Institute Sempach. Data were from the *Monitoring Häufige Brutvögel MHB* program, a common breeding bird survey (Schmid et al., 2001). The data spanned years 1999 to 2020. The survey methodology involved skilled birdwatchers conducting annual repeat transect surveys across 267 individual 1km x 1km squares laid out as a grid across Switzerland. Transect routes and squares did not change between years. Observers record all birds seen or heard during the transects. 267 unique points were surveyed over 21 years (Fig. 1C; Schmid et al., 2001).

#### United Kingdom

Data were provided in December 2021 from the British Trust for Ornithology. Data were from the *BTO/JNCC/RSPB Breeding Bird Survey* (*BBS*) and spanned years 1994 to 2015. This survey consisted of two repeat visits at the beginning and end of the breeding season of 1-km transects within an allocated 1-km square, recording all birds seen or heard (Gregory & Baillie, 1994). We summed all counts for all detected distances from the transect line for an annual count at each survey point. Between years, a stratified random sample of survey squares was selected, where stratification was representative of habitats and regions. 5,810 unique locations were surveyed over 16 years (Fig. 1E).

Within each country, we aggregated point- or transect-level count data into regional population indices. We used country-level administrative boundaries which resulted in summing our data across 16 counties in Norway, 20 counties in Sweden, 15 cantons in Switzerland, and 16 local administrative units (Nomenclature of Territorial Units for Statistics, NUTS-2) in the United Kingdom (Fig. 1). Aggregating point counts into one value for the sum of all surveyed points in a region allowed us to reduce the noise (i.e., any random fluctuation) that was present in the data and improve our ability to assess regional-level population dynamics, which was our main interest. For the United Kingdom, we took the average value of the aggregated points to account for methodological variation in the density of sample units (Link & Sauer, 2002). Small administrative units were merged to secure a minimum number of sampling locations per administrative unit (Fig. 1). From these aggregated population indices, we excluded species that were absent from at least 25% of the aggregated sites. We also excluded sites in which a species was not observed for at least 10 years of the survey duration.

Directional, temporal trends in abundance impact the strength of correlation between populations (Loreau & de Mazancourt, 2008). These directional trends can be accounted for in spatial population synchrony analyses by estimating synchrony of population growth rates instead of abundances (Loreau and de Mazancourt 2008), effectively diminishing the impacts of increasing or decreasing population abundance (Tredennick et al., 2017). Here, we calculate synchrony on both population growth rate (instantaneous rate of increase, log Nt+1/Nt) and abundance (log(*N_t_*)), but focus our interpretation of results on log population growth rate in order to consider synchrony not impacted by trends.

We classified each species along the slow-fast life history continuum using generation time as a proxy (Bird et al., 2020). Species’ generation times are defined as the average age of parents of a current cohort (IUCN, 2019) and are a common tool to distinguish species life history traits (Gaillard et al., 2005). Species-specific generation time was taken from Bird et al. (2020), which classified the worlds birds using derived generation times from proxies based on age of first reproduction, maximum longevity, and annual adult survival (Appendix 1). Where species-specific generation time was unavailable, we used generation time of the species’ next closest phylogenetic relative (2 out of 94 instances; Appendix 1).

We classified each species within each country as a resident, short-distance migrant, or long-distance migrant (Appendix 1). Migratory avian species are typically classified by the distance that they move between breeding grounds and overwintering areas (Rappole, 2013). Residents were defined as nonmigrants that made no seasonal movements outside their country of residence (Newton, 2008, Eyres et al., 2017). Short-distance migrants were defined as species that had documented nonbreeding areas within Europe, but outside the country that contained the breeding ground (Rappole, 2013). Long-distance migrants were defined as species that had documented nonbreeding areas outside of Europe (Rappole, 2013). To assign each species one of the three migration tactics (i.e. residents, short-, or long-distance migrants), we used an available avian life history trait database (Storchová & Hořák, 2018). We next confirmed country-specific species migration tactics by consulting country-specific avian information platforms [Bird Life International and the Royal Society for the Protection of Birds (UK), Swiss Ornithological Institute Swiss Breeding Bird Atlas (Knaus et al., 2020), Swedish Bird Ringing Atlas / Svensk Ringmärkningsatlas (Fransson & Hall-Karlsson, 2008), and Norwegian Bird Ringing Centre (Bakken et al., 2006)]. When country-specific avian information platforms were inconclusive, we consulted The Eurasian African Bird Migration Atlas (Franks et al., 2022) to reclassify species given their country of origin based on ringing recoveries and satellite tagging data (Kays et al., 2015, Franks et al., 2022).

### Calculating synchrony

From the aggregated abundances, we calculated the mean spatial population synchrony in two ways; either between log-transformed population growth rates (log(*N_t_* + 1 / *N_t_*)) or between log-transformed abundances (log(*N_t_*)) for each species and country separately. We log-transformed the abundance data and species’ generation times to reduce the correlation between the mean and variance. In program R (R Core Team 2020), we used a parametric Gaussian cross-correlation function to estimate synchrony between pairs of sites. Mean synchrony for each species within each country was then calculated as the mean of these estimates between pairs of sites. Distances between populations were calculated as the Euclidean distances in kilometers from the centroid coordinate of each aggregated administrative unit for each pair of sites.

Distance over which populations are sampled is known to influence estimated average spatial population synchrony (Hanski & Woiwod, 1993, Sutcliffe et al., 1996, Bjørnstad et al., 1999). This comes from the general negative relationship between population synchrony and distance between populations (Lande et al., 1999). Accordingly, for a given species, the mean synchrony would be lower if populations are sampled over large distances, compared to a smaller focal area. Given the large differences in pair-wise population distances among the four countries analyzed (e.g., max distance between aggregated points in Switzerland of 223km, max distance between aggregated points in Norway of 1,553km; Fig. 1A, Table 1), we ran all tests on mean spatial population synchrony calculated between all pairs of populations within distance thresholds 0-350km, 0-500km, 0-1,000km, and 0 – max distance interval. Statistical analyses were run separately for the four distance intervals.

**Table 1.**
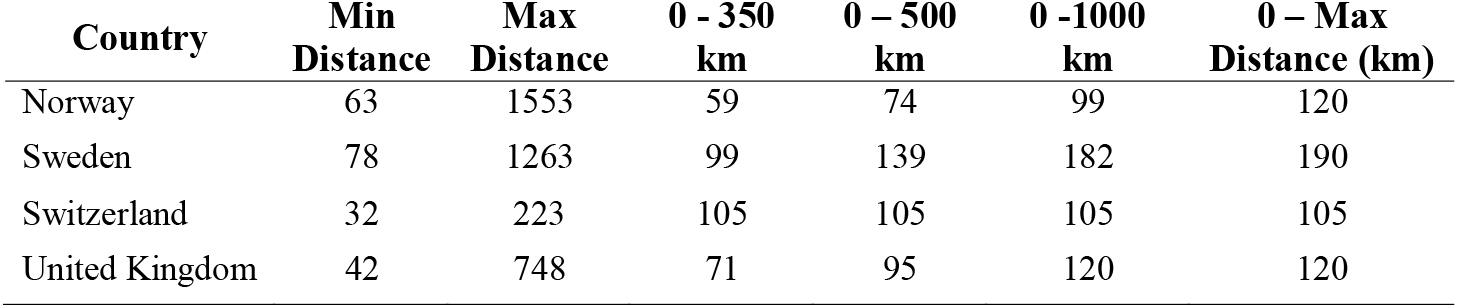
Number of paired sites for each distance interval per country. Minimum distance (min distance) calculated as the smallest distance (km) from the centroid of one aggregated administrative unit to another. Maximum distance (max distance) calculated as the largest distance (km) from the centroid of one aggregated administrative unit to another.

### Statistical analyses

To quantify the contribution of generation time and seasonal migration tactic to spatial population synchrony, we used linear mixed models. By using species as a random factor, we accounted for the nonindependence in species which was present in multiple datasets. The fixed factors in the global model included migration tactic, generation time, country, as well as all two-way interaction terms (for global model, see Table 2). We included country as a parameter to control for differences in sampling methods, survey efforts, and the variation in size of the aggregated administrative units between countries. We assumed that the environmental autocorrelation that the species experienced within countries did not differ in a meaningful way to cause species-specific differences in synchrony within each country. We included two-way interactions between country and generation time as well as country and migration tactic to test for a different effect across sampled countries for both parameters. We also included a two-way interaction between generation time and migration tactic, as we were interested in testing if different generation times were more or less sensitive to the shared environment in wintering areas. We used Akaike information criterion adjusted for small sample size (AIC_c_) to rank models (Burnham & Anderson, 2002). We assessed model uncertainty by computing simulated distributions of all parameters in the model (Knowles & Frederick, 2020). All residuals were tested for normality.

**Table 2.** Top model results for estimates of spatial population synchrony in log population growth rate and log abundance at four distance intervals (0-350km, 0-500km, 0-1000km, and 0-max distance). The inclusion of parameters of log generation time (GT), migration tactic (MT), country, and interactions between parameters designated with an “X” when present in the model. We relied upon Akaike’s Information Criterion with a small sample size correction (AIC_c_) for model selection and used Akaike model weights (AIC_c_ wt) and ΔAIC_c_ to identify the top model. Number of parameters in model indicated by column k. Top five models in each distance interval are presented. Bold models in 0-Max distance intervals were used for figures and results interpretation.

|  | Distance<br>(km) | Rank | GT | MT | Country | GT X<br>Country | GT X MT | MT X<br>Country | Log-<br>likelihood | AIC<sub>c</sub> wt | Δ AIC<sub>c</sub> | k |
| --- | --- | --- | --- | --- | --- | --- | --- | --- | --- | --- | --- | --- |
| Log Population Growth Rate | 0-350 | 1 | X | X | X |  |  |  | 180.22 | 0.39 | 0 | 9 |
|  |  | 2 | X |  | X |  |  |  | 177.14 | 0.16 | 1.77 | 7 |
|  |  | 3 | X | X | X |  |  | X | 186.03 | 0.13 | 2.14 | 15 |
|  |  | 4 | X | X | X | X |  |  | 182.39 | 0.12 | 2.43 | 12 |
|  |  | 5 | X |  | X | X |  |  | 179.48 | 0.06 | 3.71 | 10 |
|  | 0-500 | 1 | X | X | X |  |  | X | 195.29 | 0.29 | 0 | 15 |
|  |  | 2 | X | X | X |  |  |  | 188.24 | 0.24 | 0.69 | 9 |
|  |  | 3 | X |  | X |  |  |  | 185.29 | 0.11 | 1.66 | 7 |
|  |  | 4 | X | X | X | X |  | X | 197.93 | 0.11 | 2.01 | 18 |
|  |  | 5 | X | X | X | X |  |  | 190.69 | 0.079 | 2.87 | 12 |
|  | 0-1000 | 1 | X | X | X |  |  | X | 205.49 | 0.44 | 0 | 15 |
|  |  | 2 | X | X | X |  |  |  | 198.06 | 0.25 | 1.11 | 9 |
|  |  | 3 | X | X | X | X |  | X | 207.24 | 0.07 | 3.74 | 18 |
|  |  | 4 | X |  | X |  |  |  | 194.53 | 0.07 | 3.77 | 7 |
|  |  | 5 | X | X | X |  | X | X | 205.86 | 0.06 | 4.06 | 17 |
|  | 0-Max<br>distance | **1** | **X** | **X** | **X** |  |  |  | **176.13** | **0.49** | **0** | **9** |
|  |  | 2 | X | X | X |  |  | X | 182.31 | 0.24 | 1.39 | 15 |
|  |  | 3 | X | X | X |  | X |  | 176.62 | 0.08 | 3.51 | 11 |
|  |  | 4 | X | X | X |  | X | X | 183.08 | 0.05 | 4.66 | 17 |
|  |  | 5 | X |  | X | X |  |  | 177.14 | 0.05 | 4.74 | 12 |

|  | Distance<br>(km) | Rank | GT | MT | Country | GT X<br>Country | GT X MT | MT X<br>Country | Log-<br>likelihood | AIC<sub>c</sub> wt | ΔAIC<sub>c</sub> | k |
| --- | --- | --- | --- | --- | --- | --- | --- | --- | --- | --- | --- | --- |
| Log Abundance | 0-350 | 1 | X | X | X |  |  |  | 111.14 | 0.39 | 0 | 9 |
|  |  | 2 | X | X | X |  | X |  | 113.22 | 0.33 | 0.32 | 11 |
|  |  | 3 | X |  | X |  | X | X | 119.12 | 0.1 | 2.59 | 17 |
|  |  | 4 | X | X | X |  |  | X | 116.36 | 0.074 | 3.32 | 15 |
|  |  | 5 | X | X | X | X |  |  | 111.99 | 0.03 | 5.06 | 12 |
|  | 0-500 | 1 | X | X | X |  |  |  | 110.41 | 0.45 | 0 | 9 |
|  |  | 2 | X | X | X |  | X |  | 112.4 | 0.35 | 0.5 | 11 |
|  |  | 3 | X | X | X |  | X | X | 117.44 | 0.047 | 4.49 | 17 |
|  |  | 4 | X |  | X |  |  |  | 105.82 | 0.037 | 4.8 | 7 |
|  |  | 5 | X | X | X |  |  | X | 114.78 | 0.034 | 5.02 | 15 |
|  | 0-1000 | 1 | X | X | X |  |  |  | 109.71 | 0.46 | 0 | 9 |
|  |  | 2 | X | X | X |  | X |  | 111.77 | 0.38 | 0.37 | 11 |
|  |  | 3 | X | X | X | X |  |  | 110.86 | 0.05 | 4.46 | 12 |
|  |  | 4 | X | X | X | X | X |  | 112.81 | 0.03 | 5.2 | 14 |
|  |  | 5 | X |  | X |  |  |  | 104.7 | 0.03 | 5.65 | 7 |
|  | 0-Max<br>distance | 1 | X | X | X |  | X |  | 90.11 | 0.5 | 0 | 11 |
|  |  | 2 | X | X | X |  |  |  | **87.54** | **0.36** | **0.65** | **9** |
|  |  | 3 | X | X | X | X | X |  | 91.3 | 0.051 | 4.52 | 14 |
|  |  | 4 | X | X | X | X |  |  | 88.87 | 0.046 | 4.4 | 12 |
|  |  | 5 | X | X | X |  | X | X | 93.97 | 0.02 | 6.35 | 17 |

## Results

We analyzed population abundances for spatial population synchrony in 192 country-specific birds, yielding estimates of synchrony calculated for a total of 94 unique species: 36 species from Norway, 59 from Sweden, 47 from Switzerland, and 50 from the United Kingdom (Fig. 2A, Appendix 1). Most species were present in more than one country (Fig. 2A). All countries except the UK had more short-distance migrants than residents or long-distance migrants (Fig. 2B).

**Figure 2.**
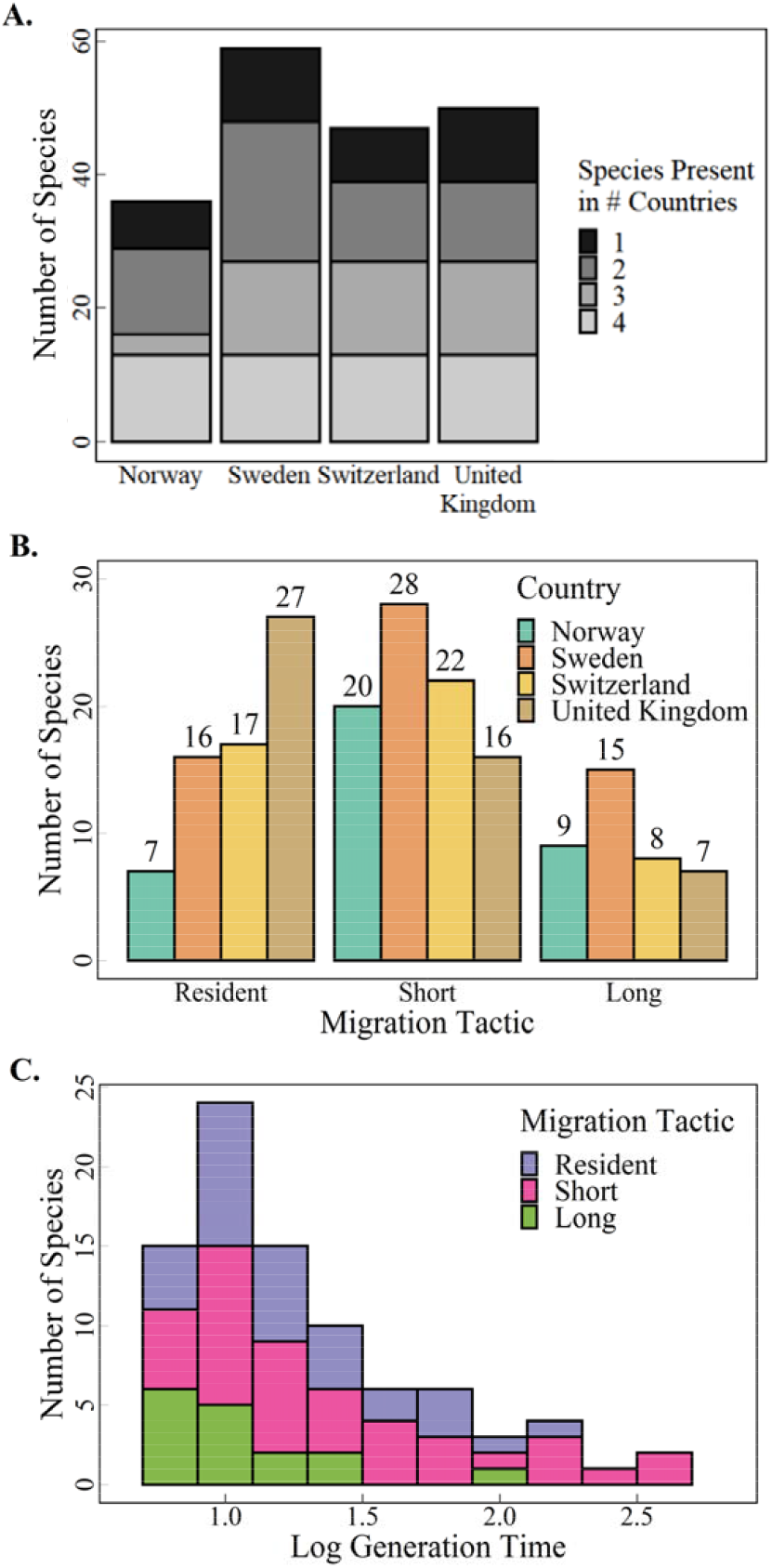
Summary of data used to estimate spatial population synchrony. (A) Number of species per country and number of species shared across multiple countries, (B) number of migration tactics per country, and (C) distribution of log generation time separated by migration tactic. Log generation time ranged from 0.53 (absolute scale: 1.69) to 2.83 (absolute scale: 16.9).

Log generation time ranged from 0.53 (absolute scale: 1.69) to 2.83 (absolute scale: 16.9; Fig. 2C). Long-distance migrants had the shortest mean log generation time (1.06), followed by resident species and short-distance migrants (1.24 and 1.30 respectively). Other life history traits associated with placement on the slow-fast life

**Figure 3.**
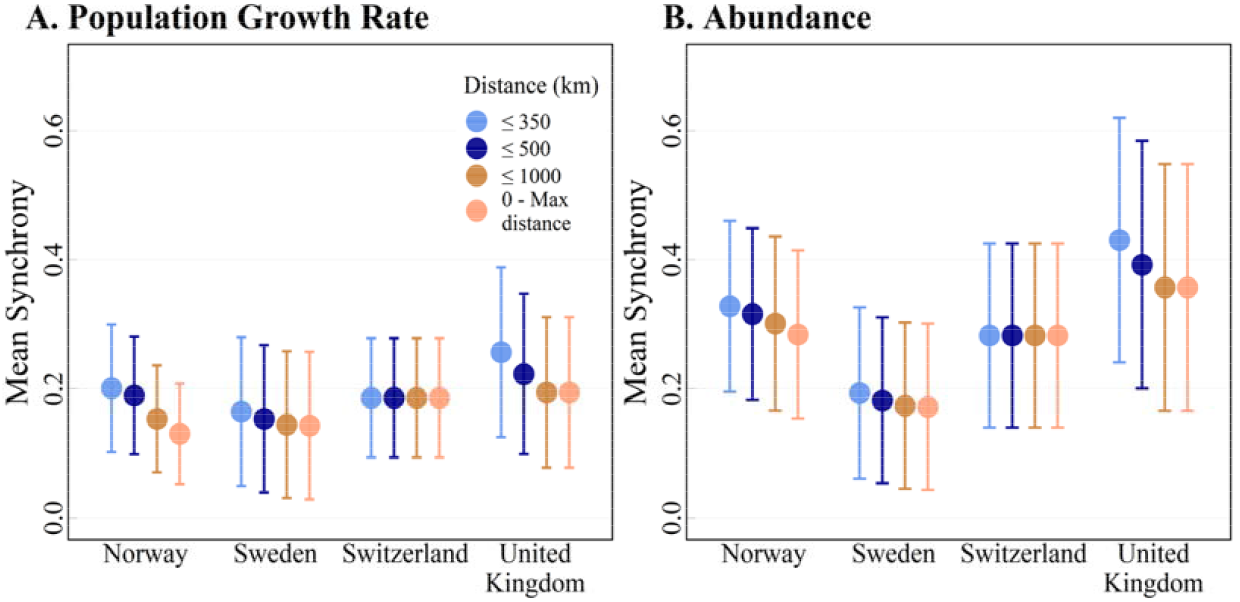
Mean synchrony (i.e., pairwise correlation in population growth rate and abundance) of all species per distance interval. Results shown for (A) log population growth rate and (B) log abundance. Number of pairs of populations per distance interval per country available in Table 1. Bars show the standard deviation.

history continuum such as survival, fecundity, and life span were highly correlated with generation time (Pearsons corr = 0.87, 0.84, 0.88 respectively; estimates for life history traits from Eyres et al., 2017, Bird et al., 2020).

Overall, mean synchrony decreased when populations at greater distances were included in analysis to estimate mean spatial population synchrony (Fig. 3). However, this relationship was weak for both growth rate (Fig. 3A, Appendix 2) and abundance (Fig. 3B, Appendix 3) and did not influence the structure of the highest ranked model, and thus the conclusions are valid over all distance classes (Table 2). Figures and results presented hereafter are generated using data from 0 – max distance intervals.

Across all distance intervals for synchrony in population growth rate, the highest ranked models included the main effects of country, migration tactic, and generation time, and in some cases an interaction between migration tactic and country (Table 2). The top two models across all distance intervals remained consistent and had similar support (ΔAIC_c_≤ 1.39 and Akaike model weights ≥ 0.24; Table 2). Parameter estimates for all top models for population growth rate across the four distance intervals were similar which suggested that our conclusions were not sensitive to the distance range at which synchrony was calculated (Appendix 4). After further exploration, the interaction between country and migration tactic evident in a top performing model in two distance classes (0-500km, 0-1000km) was driven by one bird species (*Sylvia communis*) which had notably high synchrony in population growth rate in the United Kingdom data compared with other countries and synchrony estimates (Appendix 2). There was also large uncertainty associated with the corresponding parameters for the interaction (Appendix 4, Appendix 5).

Across all distance intervals for abundance, the top performing models for synchrony included the main effects of country, migration tactic, and generation time (Table 2), and, in one case, an interaction between migration tactic and generation time (Table 2). Across all distance intervals, the top two models remained consistent and had similar support (ΔAIC_c_ ≤ 0.65 and Akaike model weights ≥ 0.33). Like the parameter estimates for population growth rate, parameter estimates for all top abundance models across the four distance intervals yielded similar parameter estimates (Appendix 4). In one distance interval, the strength of the relationship between synchrony and generation time depended on the migration tactic (Table 2, Appendix 5). This interaction appeared in only one distance interval as top model for abundance (0-max distance [km]), and there was large uncertainty associated with all of the corresponding parameters (e.g., [Short distance migrant x Log Generation Time: estimate=−0.13 SE=0.06], [Long distance migrant x Log Generation Time: estimate=−0.03 SE=0.09]).

The highest ranked models suggested that spatial population synchrony decreased with increasing generation time both for population growth rate (−0.12 [CI=-0.16 – −0.08]) and abundance (−0.14 [CI=-0.19 – −0.08], Fig. 4). Moreover, short distance migrants in general had the highest synchrony (population growth rate: 0.25, [95% confidence interval (CI)=0.19 – 0.32]; abundance: 0.48 [CI=0.39-0.57]), followed by resident species (population growth rate: 0.22 [CI=0.15 – 0.28]; abundance: 0.42 [CI=0.33-0.51]), and finally long-distance migrants (population growth rate: 0.18 [CI=0.11 – 0.24]; abundance: 0.37 [CI=0.28-0.46]). Estimates of synchrony in short-distance migrants was not significantly different from estimates of synchrony in resident species but was significantly different from estimates of synchrony in long-distance migrants (Fig. 4).

**Figure 4.**
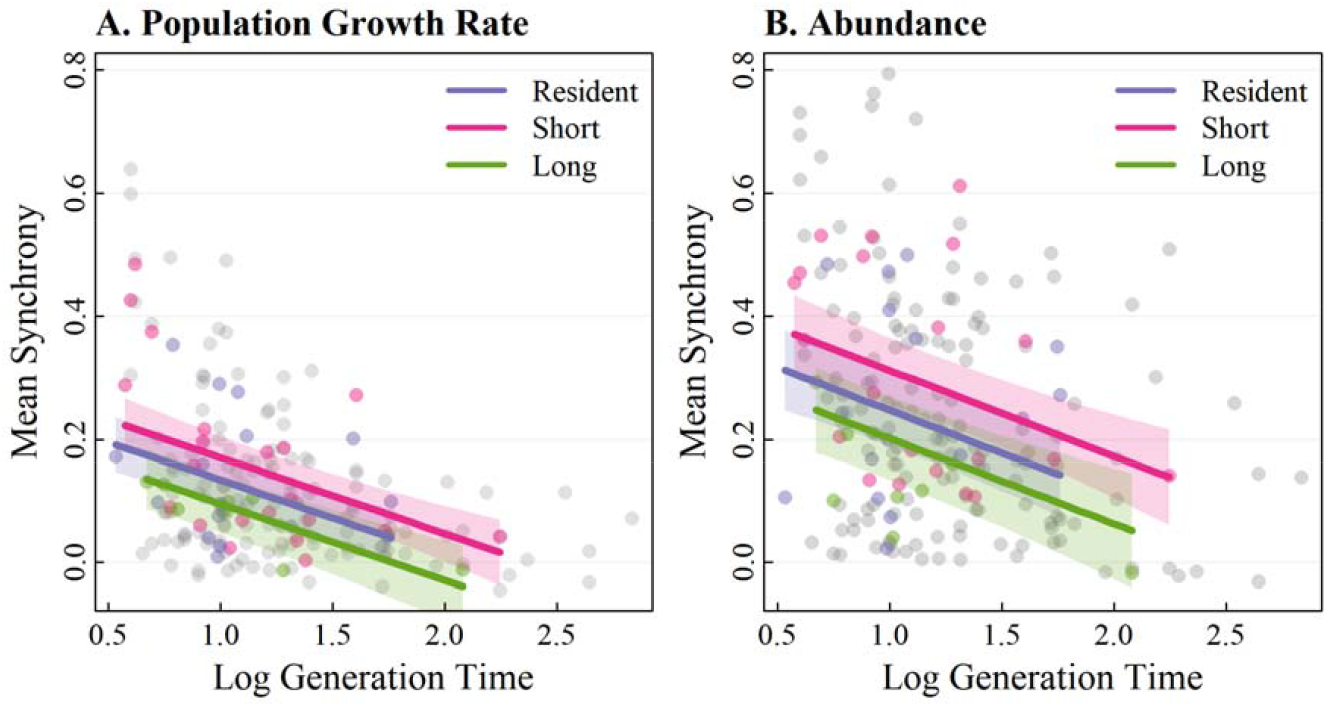
The effects of log generation time and migration tactic on mean synchrony (i.e., pairwise correlation in population growth rate and abundance) in (A) log population growth rate and (B) log abundance. Data for Switzerland in color, all other countries in grey. Slopes are predicted for Switzerland from the top performing model: Country + Migration Tactic + Log Generation Time, see Table 2. 95% confidence intervals presented as shaded colors.

Country was an important predictor of spatial population synchrony. However, there were no interactions between country and generation time or migration tactic, so the slopes and relationships between migration tactic and generation time remained the same across countries. Synchrony in growth rate was highest in the United Kingdom, followed by Switzerland, Sweden, and Norway (Appendix 4). In abundance, the highest spatial population synchrony was in the United Kingdom, followed by Norway, Switzerland, and Sweden (Appendix 4).

## Discussion

Despite the need to identify drivers of spatial population synchrony in nature, current understanding of these drivers remains more theoretical and general than species- or trait-specific. Here we make use of available long-term monitoring data to investigate synchrony across countries and species to identify life history traits that can explain why some species are more synchronized than others. Our top models confirmed that spatial population synchrony was related to species’ generation time: Species that had shorter generation times were more synchronized (Fig. 3), regardless of the spatial scale at which mean synchrony was estimated (Table 2). We also identified differences in synchrony for different migration tactics (Fig. 4). Short-distance migrants had higher synchrony in both population growth rate and abundance than long-distance migrants (Fig. 4). These results might continue to bridge a notable gap in understanding spatial population synchrony across species. Our results link known drivers of spatial population synchrony, environmental and demographic stochasticity, to species life history traits and show how these different mechanisms can be combined to understand species-specific patterns of spatial population synchrony.

We found that population synchrony was highest for species with short generation times. Theoretical and empirical examples suggest that the impact of environmental stochasticity is greater for population dynamics of species with shorter generation times (Sæther et al., 2013) and stronger density regulation, which is typically correlated with species at the fast end of the slow-fast life history continuum (Beddington & May, 1977).

Accordingly, species with shorter generation times are more sensitive to environmental stochasticity that often has a high spatial autocorrelation (Herfindal et al. 2022), and thus more synchronized than species with long generation times. At the same time, the slower dynamics of species with longer generation times can mean that fluctuations in population size have more time to spread out in space, causing synchrony over larger distances. This was found in a study of marine fish, where species with longer generation times had longer spatial scaling in synchrony, i.e., a greater distance at which spatial synchrony was below a certain value given the standard deviation, than fish with shorter generation times (Marquez et al., 2019). While spatial scaling of population synchrony has not been the focus of our current study, an interesting future question would be whether this pattern found in fish also holds for birds.

Migration is a complex phenomenon which has considerable inter- and intra-specific variation (Newton, 2008). The great diversity of migratory strategies seen in nature makes it challenging to form generalizable conclusions applicable to all migrant species. Here, we attempt to distill a complex migratory system into three generalizable categories - resident species, short-distance migrants, and long-distance migrants - to understand the influence of seasonal environments and environmental stochasticity on population synchrony. We expected to find highest synchrony in resident species because they are assumed to experience the most consistent environments and spent the most time on the breeding ground. We expected to find the lowest synchrony in long-distance migrants. Finding higher synchrony in short-distance migrants than long-distance migrants was therefore unsurprising but finding no significant differences between short-distance migrants and residents was surprising. It is possible that short-distance migrants were not significantly more synchronized than resident species because the short distance migrant species exhibited a telescopic migration strategy, where they were clustered on the wintering grounds, and thus experienced a stronger synchronizing environment on the wintering grounds [e.g., songbirds species (Beauchamp, 2011, La Sorte et al., 2016)]. It is also possible that the seasonal differences experienced by resident species reflect large seasonal differences in the scaling of environmental stochasticity on the breeding ground. In nature, there are distinct seasonal differences in environmental synchrony, particularly in terrestrial systems (Herfindal et al., 2022). This varying seasonality on the breeding grounds could have a large impact on the scaling of spatial population synchrony. A final consideration is that residents were defined by being resident within a country, but it is possible that they were altitudinal or dispersive migrants and migrated within the country borders. In these cases, variation in environment was not accounted for and could be a potential driver of the lower synchrony seen in resident birds.

As expected, long-distance migrants had the lowest spatial population synchrony. In our study, we did not investigate the driver of this lower spatial population synchrony. However, we know that long-distance migrants tend to spend the shortest amount of time on the breeding grounds before migrating across different migratory stop-over sites and wintering sites (Knaus et al. 2018). Further, the differences in sensitivity to environmental stochasticity could be driving the differences that we see between short- and long-distance migrants and residents: long-distance migrants tend to be more specialist species and are more severely affected by environmental stochasticity (Knaus et al., 2018).

Spatial scale affects estimates of spatial population synchrony (Pearson & Carroll, 1999, Dungan et al., 2002). We therefore analyzed our data at four different biologically relevant spatial scales to ensure that we captured all patterns in spatial population synchrony across local and larger, regional scales. Across all countries except Switzerland, synchrony decreased when including larger distances, but the results and support for the top models were not affected by the distance intervals. Given the large discrepancies in the range of maximum distances between countries, comparisons between countries should be done at the 350km scale because this is the maximum distance between pairs of populations in Switzerland.

Population growth rate yielded lower estimates of synchrony than abundance. This is unsurprising, as calculating synchrony on raw census data tends to reflect not only the synchronizing effect of regional environmental fluctuations, but also the synchronizing effects of common long-term trends (Koenig 1999). If trends exist, either negative or positive, there will be higher synchrony in abundance than in growth rate. There are known trends in abundance of many European bird species, particularly migratory birds (Ottvall et al., 2009, Knaus et al., 2020, Harris et al., 2022), and this directional, temporal trend in population abundance could explain why synchrony in abundance is higher than in population growth rate (Tredennick et al., 2017).

There may, however, be some biological relevancy for the weakly-supported interactions which should be considered. The interaction between generation time and migration tactic seen in the abundance model may results from differences in species traits and their responses to environmental and demographic stochasticity. For example, two species with different generation times could experience the same migratory and overwintering conditions, yet respond differently. We would expect migratory species with low sensitivity to environmental fluctuations (typically long-lived species) to be less affected by wintering ground environmental conditions than short-lived species, resulting in different effects of migration (Appendix 5). It is also possible that this interaction manifested in the abundance model set and not the population growth rate model set because of different population trends among groups of birds, which would affect synchrony in abundance but not necessarily population growth rate. Given that migratory species’ abundances are declining more than other species, estimating synchrony on abundance would pick up these trends in the data (Gilroy et al., 2016).

Further, there may be country-specific variation in synchrony across migration tactics, as seen in the population growth rate top model set. We would expect to see different synchrony for different migration tactics across countries when there is a large difference in maximum distances within each country (Norway: 1553km, Sweden: 1263km, Switzerland: 233km, UK: 748km). This large distance could be failing to uniformly capture within-country seasonal movement which could impact estimates of synchrony.

The higher spatial population synchrony we identified for European short-distance migrant species should alert managers to the susceptibility of these populations to stochastic events on shared breeding or nonbreeding grounds. Given their higher synchrony and known sensitivities to environmental stochasticity, these nonmigratory or short-distance migrants’ population dynamics are expected to be more susceptible to anthropogenic or climatically induced changes in environments. Understanding these trait-specific drivers of spatial population synchrony is important in the face of increasingly severe threats to biodiversity and could be key for successful future conservation outcomes. Further testing of the impact of life history traits on spatial population synchrony across taxa and environments is encouraged to uncover further important ecological patterns.

## Acknowledgments

The research was supported through the Research Council of Norway (Centres of Excellence funding scheme project no. 223257, and FRIPRO funding scheme project no. 276080). We thank Hans Schmid at the Swiss Ornithological Institute, John Atle Kålås at the Norwegian Institute for Nature Research, and Dario Massimino at the British Trust for Ornithology for assistance with data acquisition and advice. We thank all data collection volunteers in the Swiss breeding bird survey (MHB), the TOV-E bird survey, Swedish bird survey (standardrutterna), and the UK Breeding Bird Survey.

## Conflict of Interest

All authors have no conflicts of interest.

## Author Contributions

ECM, BBH, AML, and IH conceived the ideas and designed the methodology. ECM collated, cleaned, and formatted data. ECM and IH provided code for analysis of data. ECM led the writing of the manuscript with contributions from IH. All authors contributed critically to the drafts and gave final approval for publication.

## Data availability

Data code available in Github repository EllenCMartin/LifeHistory_SpatialPopulationSynch.

Datasets publicly available for download on GBIF:

Norway: June 2021, from GBIF DOI: https://doi.org/10.15468/6jmw2e Survey unit centroids provided by John Atle Kalas/NINA [personal communication]
Sweden: March 2021 from GBIF DOI: https://doi.org/10.15468/hd6w0r

Data available upon request to listed point of contact:

Switzerland: Swiss Ornithological Institute [data share agreement], Data from the regular territory mapping for the atlas of breeding birds 2013-2016 (Knaus, P., S. Antoniazza, S. Wechsler, J. Guélat, M. Kéry, N. Strebel & T. Sattler (2018): Swiss Breeding Bird Atlas 2013–2016. Distribution and population trends of birds in Switzerland and Liechtenstein. Swiss Ornithological Institute, Sempach. 648 p.), Point of contact: Hans Schmid.
United Kingdom: British Trust for Ornithology [data request EF1599224671889842], Point of contact: Dario Massimo.

## Notes

### Competing Interest Statement

The authors have declared no competing interest.

## References

Bakken, V., O. Runde, and E. Tjørve. 2006. Norsk Ringmerkingsatlas. Stavanger Museum, Stavanger, Norway.

Bauer, S., and B. J. Hoye. 2014. Migratory Animals Couple Biodiversity and Ecosystem Functioning Worldwide. Science 344:1242552

Beauchamp, G. 2011. Long-distance migrating species of birds travel in larger groups. Biology Letters 7:692–694.

Beddington, J. R., and R. M. May. 1977. Harvesting natural populations in a randomly fluctuating environment. Science 80:463–465.

Bird, J. P., R. Martin, H. R. Akçakaya, J. Gilroy, I. J. Burfield, S. T. Garnett, A. Symes, J. Taylor, Ç. H. Şekercioğlu, and S. H. M. Butchart. 2020. Generation lengths of the world’s birds and their implications for extinction risk. Conservation Biology 34:1252–1261.

Bjørkvoll, E., V. Grøtan, A. Sondre, B.-E. Sæther, E. Steinar, and R. Aanes. 2012. Stochastic Population Dynamics and Life-History Variation in Marine Fish Species. American Naturalist 180:372–387.

Bjørnstad, O. N., R. A. Ims, and X. Lambin 1999. Spatial population dynamics: analyzing patterns and processes of population synchrony. Trends in Ecology & Evolution 14:427–432.

Bogdanova, M. I., F. Daunt, M. Newell, R. A. Phillips, M. P. Harris, and S. Wanless. 2011. Seasonal interactions in the black legged kittiwake, Rissa tridactyla: links between breeding performance and winter distribution. Proceedings of the Royal Society B 278:2412–2418.

Burnham, K. P., and D. R. Anderson. 2002. Model selection and multi-model inference: a practical information-theoretic approach. 2 edition. Springer, New York, USA.

Chevalier, M., P. Laffaille, and G. Grenouillet. 2014. Spatial synchrony in stream fish populations: influence of species traits. Ecography 37:960–968.

Dungan, J. L., J. N. Perry, M. R. T. Dale, P. Legendre, S. Citron-Pousty, M. J. Fortin, A. Jakomulska, M. Miriti, and M. S. Rosenberg. 2002. A balanced view of scale in spatial statistical analysis. Ecography 25:626–640.

Ellis, J., and D. C. Schneider. 2008. Spatial and temporal scaling in benthic ecology. Journal of Experimental Marine Biology and Ecology 366:92–98.

Engen, S., R. Lande, and B.-E. Sæther. 2002. Migration and Spatiotemporal Variation in Population Dynamics in a Heterogeneous Environment. Ecology 83:570–579.

Engen, S., and B.-E. Sæther. 2005. Generalizations of the Moran effect explaining spatial synchrony in population fluctuations. American Naturalist 166:603–612.

Engen, S., and B.-E. Sæther. 2016. Spatial synchrony in population dynamics: The effects of demographic stochasticity and density regulation with a spatial scale. Mathematical Biosciences 274:17–24.

Eyres, A., K. Bohning-Gaese, and S. A. Fritz. 2017. Quantification of climatic niches in birds: adding the temporal dimension. Journal of Avian Biology 48:1517–1531.

Ferguson, S. H., and S. Larivière. 2002. Can comparing life histories help conserve carnivores? Animal Conservation 5:1–12.

Franks, S., W. Fiedler, J. Arizaga, F. Jiguet, B. Nikolov, H. van der Jeugd, Ambrosini, R., Aizpurua, O., Bairlein, F., J. Clark, N. Fattorini, Hammond, M., Higgins, D., H. Levering, Skellorn, W., F. Spina, K. Thorup, J. Walker, Woodward, I., and S. R. Baillie. 2022. Online Atlas of the movements of European bird populations. EURING/CMS.

Fransson, T., and S. Hall-Karlsson. 2008. Svensk Ringmärkningsatlas: Swedish Bird Ringing Atlas, Volume 3, Passerines. Naturhistoriska riksmuseet.

Gaillard, J. M., N. G. Yoccoz, J. D. Lebreton, C. Bonenfant, S. Devillard, A. Loison, D. Pontier, and D. Allaine. 2005. Generation time: A reliable metric to measure life-history variation among mammalian populations. American Naturalist 166:119–123.

Gilroy, J. J., J. A. Gill, S. H. M. Butchart, V. R. Jones, and A. M. A. Franco. 2016. Migratory diversity predicts population declines in birds. Ecology Letters 19:308–317.

Gregory, R. D., and S. R. Baillie. 1994. Evaluation of sampling strategies for 1km squares for inclusion in the Breeding Bird Survey. British Trust for Ornithology, Thetford.

Hansen, B. B., Å. Ø. Pedersen, B. Peeters, M. Le Moullec, S. D. Albon, I. Herfindal, B.-E. Sæther, V. Grøtan, and R. Aanes. 2019. Spatial heterogeneity in climate change effects decouples the long-term dynamics of wild reindeer populations in the high Arctic. Global Change Biology 25:3656–3668.

Hanski, I., T. Pakkala, M. Kuussaari, and G. C. Lei. 1995. Metapopulation Persistence of an Endangered Butterfly in a Fragmented Landscape. Oikos 72:21–28.

Hanski, I., and I. P. Woiwod. 1993. Spatial Synchrony in the Dynamics of Moth and Aphid Populations. Journal of Animal Ecology 62:656–668.

Harris, S. J., D. Massimino, D. E. Balmer, L. Kelly, D. G. Noble, J. W. Pearce-Higgins, P. Woodcock, S. Wotton, and S. Gillings. 2022. The Breeding Bird Survey 2021. British Trust for Ornithology, Thetford.

Harrison, X. A., J. D. Blount, R. Inger, D. R. Norris, and S. Bearhop. 2010. Carry-over effects as drivers of fitness differences in animals. Journal of Animal Ecology 80:4–18.

Heino, M., V. Kaitala, E. Ranta, and J. Lindstrom. 1997. Synchronous dynamics and rates of extinction in spatially structured populations. Proceedings of the Royal Society B-Biological Sciences 264:481–486.

Herfindal, I., S. Aanes, R. Benestad, A. G. Finstad, A. Salthaug, N. C. Stenseth, and B.-E. Saether. 2022. Spatiotemporal variation in climatic conditions across ecosystems. Climate Research 86:9–19.

Ims, R. A., and H. P. Andreassen. 2000. Spatial synchronization of vole population dynamics by predatory birds. Nature 408:194–196.

IUCN. 2019. Guide-lines for the IUCN Red List categories and criteria. International Union for the Conservation of Naure, Gland, Switzerland.

Jones, J., P. J. Doran, and R. T. Holmes. 2007. Spatial scaling of avian population dynamics: Population abundance, growth rate, and variability. Ecology 88:2505–2515.

Kålås, J. A., I. J. Øien, B. Stokke, and R. Vang. 2022. TOV-E Bird monitoring sampling data. in N. I. f. N. Research, editor.

Kays, R., M. C. Crofoot, W. Jetz, and M. Wilkelski. 2015. Terrestrial animal tracking as an eye on life and planet. Science 348:aaa2478.

Kendall, B. E., O. N. Bjørnstad, J. Bascompte, T. H. Keitt, and W. F. Fagan. 2000. Dispersal, environmental correlation, and spatial synchrony in population dynamics. American Naturalist 155:628–636.

Knaus, P., S. Antoniazza, S. Wechsler, J. Guélat, M. Kéry, N. Strebel, and T. Sattler. 2018. Swiss Breeding Bird Atlas 2013–2016. Distribution and population trends of birds in Switzerland and Liechtenstein. Swiss Ornithological Institute, Sempach.

Knaus, P., T. Sattler, H. Schmid, N. Strebel, and B. Volet. 2020. The State of Birds in Switzerland. Swiss Ornithological Institute, Sempach.

Knowles, J. E., and C. Frederick. 2020. merTools: Tools for Analyzing Mixed Effect Regression Models https://CRAN.R-project.org/package=merTools.

Koenig, W. D. 2006. Spatial synchrony of monarch butterflies. American Midland Naturalist 155:39–49.

Koenig, W. D., and A. M. Liebhold. 2016. Temporally increasing spatial synchrony of North American temperature and bird populations. Nature Climate Change 6:614–617.

La Sorte, F. A., D. Fink, W. M. Hochachka, and S. Kelling. 2016. Convergence of broad-scale migration strategies in terrestrial birds. Proceedings of the Royal Society B 283.

Lande, R., S. Engen, and B.-E. Sæther. 1999. Spatial scale of population synchrony: Environmental correlation versus dispersal and density regulation. American Naturalist 154:271–281.

Lande, R., S. Engen, and B.-E. Sæther. 2003. Stochastic population dynamics in ecology and conservation. Oxford University Press, New York.

Lande, R., S. Engen, B.-E. Sæther, F. Filli, E. Matthysen, and H. Weimerskirch. 2002. Estimating density dependence from population time series using demographic theory and life-history data. American Naturalist 159:321–337.

Lindström, Å., and M. Green. 2021. Swedish Bird Survey: Fixed routes (Standardrutterna). in L. U. Department of Biology, editor.

Link, W. A., and J. R. Sauer. 2002. A hierarchical analysis of population change with application to cerulean warblers. Ecology 83:2832–2840.

Loreau, M., and C. de Mazancourt. 2008. Species Synchrony and Its Drivers: Neutral and Nonneutral Community Dynamics in Fluctuating Environments. American Naturalist 172.

MacArthur, R. H., and E. O. Wilson. 1967. The theory of island biogeography. Princeton University Press, Princeton, N.J.

Marquez, J. E., A. M. Lee, S. Aanes, S. Engen, I. Herfindal, A. Salthaug, and B.-E. Sæther. 2019. Spatial scaling of population synchrony in marine fish depends on their life history. Ecology Letters 22:1787–1796.

Moran, P. A. P. 1953. The Statistical Analysis of the Canadian Lynx Cycle: Structure and Prediction. Australian Journal of Zoology 1:163–173.

Myrberget, S. 1973. Geographical Synchronism of Cycles of Small Rodents in Norway. Oikos 24: 220–224

Newton, I. 2008. The migration ecology of birds. Academic Press.

Oli, M. K. 2004. The fast–slow continuum and mammalian life-history patterns: an empirical evaluation. Basic and Applied Ecology 5:449–463.

Ottvall, R., L. Edenius, J. Elmberg, H. Engström, M. Green, N. Holmqvist, Å. Lindström, T. Pärt, and M. Tjernberg. 2009. Population trends for Swedish breeding birds. Ornis Svecica 19:117–192.

Paradis, E., S. R. Baillie, W. J. Sutherland, and R. D. Gregory. 1999. Dispersal and spatial scale affect synchrony in spatial population dynamics. Ecology Letters 2:114–120.

Pearson, D. L., and S. S. Carroll. 1999. The influence of spatial scale on cross-taxon congruence patterns and prediction accuracy of species richness. Journal of Biogeography 26:1079–1090.

Raimondo, S., A. M. Liebhold, J. S. Strazanac, and L. Butler. 2004. Population synchrony within and among Lepidoptera species in relation to weather, phylogeny, and larval phenology. Ecological Entomology 29:96–105.

Ranta, E., V. Kaitala, J. Lindstrom, and H. Linden. 1995. Synchrony in Population-Dynamics. Proceedings of the Royal Society B-Biological Sciences 262:113–118.

Rappole, J. H. 2013. The avian migrant: The biology of bird migration. Columbia University Press, New York.

Sæther, B.-E. 1997. Environmental stochasticity and population dynamics of large herbivores: A search for mechanisms. Trends in Ecology & Evolution 12:143–149.

Sæther, B.-E., and Ø. Bakke. 2000. Avian life history variation and contribution of demographic traits to the population growth rate. Ecology 81:642–653.

Sæther, B.-E., T. Coulson, V. Grøtan, S. Engen, R. Altwegg, K. B. Armitage, C. Barbraud, P. H. Becker, D. T. Blumstein, F. S. Dobson, M. Festa-Bianchet, J. M. Gaillard, A. Jenkins, C. Jones, M. A. Nicoll, K. Norris, M. K. Oli, A. Ozgul, and H. Weimerskirch. 2013. How life history influences population dynamics in fluctuating environments. American Naturalist 182:743–759.

Sæther, B.-E., S. Engen, V. Grøtan, W. Fiedler, E. Matthysen, M. E. Visser, J. Wright, A. P. Moller, F. Adriaensen, H. Van Balen, D. Balmer, M. C. Mainwaring, R. H. McCleery, M. Pampus, and W. Winkel. 2007. The extended Moran effect and large-scale synchronous fluctuations in the size of great tit and blue tit populations. Journal of Animal Ecology 76:315–325.

Sæther, B.-E., R. Lande, S. Engen, H. Weimerskirch, M. Lillegård, R. Altwegg, P. H. Becker, T. Bregnballe, J. E. Brommer, R. H. McCleery, J. Merilä, E. Nyholm, W. Rendell, R. R. Robertson, P. Tryjanowski, and M. E. Visser. 2005. Generation time and temporal scaling of bird population dynamics. Nature 436:99–102.

Schmid, H., M. Burkhardt, V. Keller, P. Knaus, B. Volet, and N. Zbinden. 2001. The development of the bird world in Switzerland/L’évolution de l’avifaune en Suisse. Sempach 1.

Selonen, V., S. Helle, T. Laaksonen, M. P. Ahola, E. Lehikoinen, and T. Eeva. 2021. Identifying the paths of climate effects on population dynamics: dynamic and multilevel structural equation model around the annual cycle. Oecologia 195:525–538.

Somveille, M., R. A. Bay, T. B. Smith, P. P. Marra, and K. C. Ruegg. 2021. A general theory of avian migratory connectivity. Ecology Letters 24:1848–1858.

Stearns, S. C. 1999. The Evolution of Life Histories. Oxford University Press, New York.

Storchová, L., and D. Hořák. 2018. Life-history characteristics of European birds. Global Ecology and Biogeography 27:400–406.

Sutcliffe, O. L., C. D. Thomas, and D. Moss. 1996. Spatial synchrony and asynchrony in butterfly population dynamics. Journal of Animal Ecology 65:85–95.

Swanson, B. J., and D. R. Johnson. 1999. Distinguishing Causes of Intraspecific Synchrony in Population Dynamics. Oikos 86: 265–274.

Tedesco, P., and B. Hugueny. 2006. Life history strategies affect climate based spatial synchrony in population dynamics of West African freshwater fishes. Oikos 115:117–127.

Tredennick, A. T., G. Hooker, S. P. Ellner, and P. B. Adler. 2017. A practical guide to selecting models for exploration, inference, and prediction in ecology. Ecology 106:e03336.

Webster, M. S., P. P. Marra, S. M. Haig, S. Bensch, and R. T. Holmes. 2002. Links between worlds: unraveling migratory connectivity. Trends in Trends in Ecology & Evolution 17:76–83.

