## Supplemental Files for "Generation time and seasonal migration explain variation in spatial population synchrony across European bird species"

### **Supplementary material**

Appendix 1. Bird species in analysis and corresponding migration tactic (resident, short-distance migrant [short], or long-distance migrant [long]) indicated in country column where species was present and analyzed (Norway, Sweden, Switzerland, and United Kingdom). Species noted by asterisk (*) had different migration tactics between two or more countries. Generation time presented from Bird et al. 2020. Generation times are defined as the average age of parents of the current cohort. Asterisk in generation time column indicates species for which generation time was unavailable; value given is from closest phylogenetic relative. Totals of residents, short-distance migrants, and long-distance migrants per country given at bottom of table.

Appendix 2. Estimated mean spatial population synchrony in log population growth rate for each species by country. *NA* indicates that the species was not included in the country’s data.

Appendix 3. Estimated mean spatial population synchrony in log abundance for each species by country. *NA* indicates that the species was not in the country associated with the column. Species names in Latin and English common names provided.

Appendix 4. Parameter estimates and standard errors (parentheses) for all chosen models across all distance intervals for log population growth rate (A) and log abundance (B). Max distance varied by country, for max distance values see Table 2.

Appendix 5. Interaction terms between migration tactic and country (log population growth rate) or migration tactic and generation time (log abundance) appeared in the top model. Mean synchrony is estimated from the log population growth rate top model (Country * Migration Tactic + Log Generation Time) and 0-max distance interval log abundance top model (Country + Migration Tactic * Log Generation Time). 95% confidence intervals are presented as shaded colors.

### Appendix 1. Bird species in analysis and corresponding migration tactic (resident, short-distance migrant [short], or long-distance migrant [long]) indicated in country column where species was present and analyzed (Norway, Sweden, Switzerland, and United Kingdom). Species noted by asterisk (*) had different migration tactics between two or more countries. Generation time presented from Bird et al. 2020. Generation times are defined as the average age of parents of the current cohort. Asterisk in generation time column indicates species for which generation time was unavailable; value given is from closest phylogenetic relative. Totals of residents, short-distance migrants, and long-distance migrants per country given at bottom of table.

|  | **Country** | | | |  |
| --- | --- | --- | --- | --- | --- |
| **Species** | **Norway** | **Sweden** | **Switzerland** | **United Kingdom** | **Generation Time** |
| Acanthis flammea | Short |  |  |  | 2.59 |
| Aegithalos caudatus |  |  |  | Resident | 2.39 |
| Alauda arvensis* |  |  | Short | Resident | 2.83 |
| Anas platyrhynchos |  | Short |  | Short | 4.78 |
| Anthus pratensis | Short |  |  | Short | 2.17 |
| Anthus spinoletta |  |  | Short |  | 2.17 |
| Anthus trivialis | Long | Long | Long |  | 2.11 |
| Apus apus |  | Long | Long | Long | 8.01 |
| Ardea cinerea |  |  |  | Resident | 8.88 |
| Branta canadensis |  | Short |  |  | 9.45 |
| Bucephala clangula |  | Short |  |  | 7.12 |
| Buteo buteo* |  |  | Short | Resident | 9.45 |
| Carduelis cannabina |  |  | Short | Short | 2.20* |
| Carduelis carduelis* |  |  | Short | Resident | 2.53 |
| Certhia brachydactyla |  |  | Resident |  | 1.70 |
| Certhia familiaris |  | Resident | Resident |  | 2.05 |
| Chloris chloris* | Short | Short | Resident | Short | 2.71 |
| Chroicocephalus ridibundus |  | Short |  |  | 9.85 |
| Columba livia |  |  |  | Resident | 3.99 |
| Columba oenas |  |  |  | Resident | 3.36 |
| Columba palumbus* | Short | Short | Short | Resident | 3.72 |
| Corvus corax | Resident | Resident |  |  | 7.46 |
| Corvus corone | Resident | Resident | Resident | Resident | 5.72 |
| Corvus frugilegus |  |  |  | Resident | 5.59 |
| Corvus monedula* |  | Short |  | Resident | 5.57 |
| Cuculus canorus | Long | Long | Long | Long | 2.76 |
| Cyanistes caeruleus |  | Resident | Resident | Resident | 2.93 |
| Delichon urbicum |  | Long |  | Long | 2.92 |
| Dendrocopos major |  | Resident | Resident | Resident | 2.70 |
| Dryocopus martius |  | Resident |  |  | 4.12 |
| Emberiza citronella* | Resident | Short | Short | Resident | 2.77 |
| Emberiza schoeniclus | Short | Short |  |  | 2.46 |
| Erithacus rubecula* | Short | Short | Short | Resident | 3.60 |
| Falco tinnunculus |  |  |  | Resident | 4.08 |
| Ficedula hypoleuca | Long | Long |  |  | 4.12 |
| Fringilla coelebs | Short | Short | Short | Short | 4.98 |
| Fringilla montifringilla | Short |  |  |  | 2.99 |
| Gallinago gallinago | Short |  |  |  | 3.57 |
| Gallinula chloropus |  |  |  | Resident | 3.57 |
| Garrulus glandarius |  | Resident | Resident | Resident | 4.91 |
| Grus grus |  | Short |  |  | 17.03 |
| Hirundo rustica |  | Long | Long | Long | 3.13 |
| Lagopus lagopus | Resident |  |  |  | 2.33 |
| Lagopus muta | Resident |  |  |  | 2.99 |
| Larus argentatus |  | Short |  | Short | 14.07 |
| Larus canus |  | Short |  |  | 10.67 |
| Larus fuscus |  |  |  | Short | 12.62 |
| Lophophanes cristatus |  | Resident | Resident |  | 2.50 |
| Loxia curvirostra |  | Short |  |  | 3.16 |
| Lyrurus tetrix | Resident | Resident |  |  | 3.37 |
| Motacilla alba | Long | Long | Long | Long | 2.81 |
| Muscicapa striata | Long | Long |  |  | 2.55 |
| Oenanthe oenanthe | Long |  | Long |  | 2.25 |
| Parus major | Resident | Resident | Resident | Resident | 3.05 |
| Passer domesticus |  | Resident | Resident | Resident | 3.73 |
| Passer montanus |  | Resident | Resident |  | 2.72 |
| Periparus ater |  | Resident | Resident | Resident | 2.20 |
| Phasianus colchicus |  |  |  | Resident | 4.77 |
| Phoenicurus ochruros |  |  | Short |  | 2.41 |
| Phoenicurus phoenicurus | Long | Long |  |  | 2.31 |
| Phylloscopus bonelli |  |  | Long |  | 1.95* |
| Phylloscopus collybita | Short |  | Short | Short | 2.00 |
| Phylloscopus sibilatrix |  | Long |  |  | 2.31 |
| Phylloscopus trochilus | Long | Long |  | Long | 2.53 |
| Pica pica |  | Resident | Resident | Resident | 5.81 |
| Picus viridis |  |  |  | Resident | 3.06 |
| Pluvialis apricaria | Long |  |  |  | 4.45 |
| Poecile montanus | Resident | Resident |  |  | 2.47 |
| Poecile palustris |  |  | Resident |  | 2.57 |
| Prunella collaris |  |  | Resident |  | 2.68 |
| Prunella modularis | Short | Short | Short | Short | 3.81 |
| Pyrrhula pyrrhula* |  |  | Short | Resident | 3.35 |
| Regulus ignicapilla |  |  | Short |  | 1.78 |
| Regulus regulus* |  | Short | Short | Resident | 1.85 |
| Saxicola rubetra |  | Long |  |  | 1.91 |
| Serinus serinus |  |  | Short |  | 2.48 |
| Sitta europaea |  | Resident | Resident |  | 2.69 |
| Spinus spinus | Short | Short |  |  | 2.78 |
| Streptopelia decaocto |  |  |  | Resident | 3.46 |
| Sturnus vulgaris |  | Short | Short | Short | 5.65 |
| Sylvia atricapilla |  | Short | Short | Short | 2.51 |
| Sylvia borin |  | Long | Long |  | 3.59 |
| Sylvia communis |  | Long |  | Long | 2.17 |
| Sylvia curruca |  | Long |  |  | 2.14 |
| Tringa ochropus |  | Short |  |  | 4.75 |
| Tringa tetanus | Short |  |  |  | 4.75 |
| Troglodytes troglodytes | Short | Short | Short | Short | 1.82 |
| Turdus iliacus | Short | Short |  |  | 3.53 |
| Turdus merula | Short | Short | Short | Short | 4.03 |
| Turdus philomelos | Short | Short | Short | Short | 3.37 |
| Turdus pilaris | Short | Short |  |  | 3.43 |
| Turdus torquatus | Short |  | Short |  | 2.99 |
| Turdus viscivorus |  | Short | Short | Short | 3.97 |
| Vanellus vanellus |  | Short |  | Short | 6.19 |
| Total Species: | 36 | 59 | 47 | 50 |  |
| Residents | 7 | 16 | 17 | 27 |  |
| Short-Distance Migrants | 20 | 28 | 22 | 16 |  |
| Long-Distance Migrants | 9 | 15 | 8 | 7 |  |

### Appendix 2. Estimated mean spatial population synchrony in log population growth rate for each species by country. *NA* indicates that the species was not included in the country’s data.

| **Species** | **Common Name** | **Norway** | **Sweden** | **Switzerland** | **United Kingdom** |
| --- | --- | --- | --- | --- | --- |
| *Acanthis flammea* | Common Redpoll | 0.36 | NA | NA | NA |
| *Aegithalos caudatus* | Long-tailed Tit | NA | NA | NA | 0.18 |
| *Alauda arvensis* | Eurasian Skylark | NA | NA | 0.02 | 0.18 |
| *Anas platyrhynchos* | Mallard | NA | 0.02 | NA | 0.02 |
| *Anthus pratensis* | Meadow Pipit | 0.08 | NA | NA | -0.005 |
| *Anthus spinoletta* | Water Pipit | NA | NA | 0.09 | NA |
| *Anthus trivialis* | Tree Pipit | 0.08 | 0.04 | 0.13 | NA |
| *Apus apus* | Common Swift | NA | -0.003 | -0.01 | 0.05 |
| *Ardea cinerea* | Grey Heron | NA | NA | NA | 0.11 |
| *Branta canadensis* | Canada Goose | NA | -0.05 | NA | NA |
| *Bucephala clangula* | Common Goldeneye | NA | -0.01 | NA | NA |
| *Buteo buteo* | Eurasian Buzzard | NA | NA | 0.04 | 0.04 |
| *Carduelis cannabina* | Common Linnet | NA | NA | NA | 0.15 |
| *Carduelis carduelis* | European Goldfinch | NA | NA | 0.22 | 0.19 |
| *Certhia brachydactyla* | Short-toed Treecreeper | NA | NA | 0.17 | NA |
| *Certhia familiaris* | Eurasian Treecreeper | NA | 0.20 | 0.10 | NA |
| *Chloris chloris* | European Greenfinch | 0.10 | 0.04 | 0.03 | 0.22 |
| *Chroicocephalus ridibundus* | Black-headed Gull | NA | -0.02 | NA | NA |
| *Columba livia* | Rock Dove | NA | NA | NA | 0.05 |
| *Columba oenas* | Stock Dove | NA | NA | NA | 0.02 |
| *Columba palumbus* | Common Woodpigeon | 0.13 | 0.03 | 0.10 | 0.11 |
| *Corvus corax* | Common Raven | 0.04 | 0.13 | NA | NA |
| *Corvus corone* | Carrion Crow | 0.04 | 0.04 | 0.04 | 0.02 |
| *Corvus frugilegus* | Rook | NA | NA | NA | -0.04 |
| *Corvus monedula* | Western Jackdaw | NA | 0.03 | NA | 0.03 |
| *Cuculus canorus* | Common Cuckoo | 0.12 | 0.06 | 0.08 | 0.12 |
| *Cyanistes caeruleus* | Eurasian Blue Tit | NA | 0.10 | 0.28 | 0.31 |
| *Delichon urbicum* | Common House Martin | NA | -0.01 | NA | 0.15 |
| *Dendrocopos major* | Great Spotted Woodpecker | NA | 0.20 | 0.29 | 0.03 |
| *Dryocopus martius* | Black Woodpecker | NA | 0.12 | NA | NA |
| *Emberiza citrinella* | Yellowhammer | 0.02 | 0.06 | NA | 0.08 |
| *Emberiza schoeniclus* | Common Reed Bunting | -0.01 | -0.02 | NA | NA |
| *Erithacus rubecula* | European Robin | 0.26 | 0.23 | 0.19 | 0.30 |
| *Falco tinnunculus* | Common Kestrel | NA | NA | NA | 0.31 |
| *Ficedula hypoleuca* | European Pied Flycatcher | 0.04 | 0.03 | NA | NA |
| *Fringilla coelebs* | Common Chaffinch | 0.15 | 0.15 | 0.27 | 0.12 |
| *Fringilla montifringilla* | Brambling | 0.01 | NA | NA | NA |
| *Gallinago gallinago* | Common Snipe | 0.03 | NA | NA | NA |
| *Gallinula chloropus* | Common Moorhen | NA | NA | NA | 0.06 |
| *Garrulus glandarius* | Eurasian Jay | NA | 0.09 | 0.20 | 0.09 |
| *Grus grus* | Common Crane | NA | 0.07 | NA | NA |
| *Hirundo rustica* | Barn Swallow | NA | -0.01 | 0.11 | 0.15 |
| *Lagopus lagopus* | Willow Ptarmigan | 0.15 | NA | NA | NA |
| *Lagopus muta* | Rock Ptarmigan | 0.13 | NA | NA | NA |
| *Larus argentatus* | European Herring Gull | NA | -0.03 | NA | 0.02 |
| *Larus canus* | Common Gull | NA | 0.01 | NA | NA |
| *Larus fuscus* | Lesser Black-backed Gull | NA | NA | NA | 0.12 |
| *Lophophanes cristatus* | Crested Tit | NA | 0.25 | 0.16 | NA |
| *Loxia curvirostra* | Red Crossbill | NA | 0.16 | NA | NA |
| *Lyrurus tetrix* | Black Grouse | -0.01 | 0.17 | NA | NA |
| *Motacilla alba* | White Wagtail | 0.08 | -0.01 | 0.01 | 0.16 |
| *Muscicapa striata* | Spotted Flycatcher | 0.03 | 0.04 | NA | NA |
| *Oenanthe oenanthe* | Northern Wheatear | 0.03 | NA | 0.09 | NA |
| *Parus major* | Great Tit | 0.06 | 0.03 | 0.21 | 0.17 |
| *Passer domesticus* | House Sparrow | NA | -0.01 | 0.06 | 0.17 |
| *Passer montanus* | Eurasian Tree Sparrow | NA | 0.09 | 0.10 | NA |
| *Periparus ater* | Coal Tit | NA | 0.11 | 0.35 | 0.16 |
| *Phasianus colchicus* | Common Pheasant | NA | NA | NA | 0.05 |
| *Phoenicurus ochruros* | Black Redstart | NA | NA | 0.16 | NA |
| *Phoenicurus phoenicurus* | Common Redstart | 0.04 | 0.04 | NA | NA |
| *Phylloscopus bonelli* | Western Bonelli's Warbler | NA | NA | 0.13 | NA |
| *Phylloscopus collybita* | Common Chiffchaff | 0.03 | NA | 0.38 | 0.39 |
| *Phylloscopus sibilatrix* | Wood Warbler | NA | 0.05 | NA | NA |
| *Phylloscopus trochilus* | Willow Warbler | 0.11 | 0.13 | NA | 0.30 |
| *Pica pica* | Eurasian Magpie | NA | 0.06 | 0.10 | 0.08 |
| *Picus viridis* | European Green Woodpecker | NA | NA | NA | 0.09 |
| *Pluvialis apricaria* | European Golden Plover | 0.01 | NA | NA | NA |
| *Poecile montanus* | Willow Tit | 0.05 | 0.11 | NA | NA |
| *Poecile palustris* | Marsh Tit | NA | NA | 0.04 | NA |
| *Prunella collaris* | Alpine Accentor | NA | NA | 0.01 | NA |
| *Prunella modularis* | Dunnock | 0.04 | 0.10 | 0.04 | 0.10 |
| *Pyrrhula pyrrhula* | Eurasian Bullfinch | NA | NA | 0.18 | 0.24 |
| *Regulus ignicapilla* | Common Firecrest | NA | NA | 0.29 | NA |
| *Regulus regulus* | Goldcrest | NA | 0.49 | 0.48 | 0.42 |
| *Saxicola rubetra* | Whinchat | NA | 0.02 | NA | NA |
| *Serinus serinus* | European Serin | NA | NA | 0.06 | NA |
| *Sitta europaea* | Eurasian Nuthatch | NA | 0.38 | 0.08 | NA |
| *Spinus spinus* | Eurasian Siskin | 0.37 | 0.49 | NA | NA |
| *Streptopelia decaocto* | Eurasian Collared Dove | NA | NA | NA | 0.01 |
| *Sturnus vulgaris* | Common Starling | NA | 0.01 | 0.05 | 0.13 |
| *Sylvia atricapilla* | Eurasian Blackcap | NA | 0.29 | 0.20 | 0.30 |
| *Sylvia borin* | Garden Warbler | NA | 0.19 | -0.01 | NA |
| *Sylvia communis* | Common Whitethroat | NA | 0.17 | NA | 0.50 |
| *Sylvia curruca* | Lesser Whitethroat | NA | 0.08 | NA | NA |
| *Tringa ochropus* | Green Sandpiper | NA | 0.07 | NA | NA |
| *Tringa totanus* | Common Redshank | 0.02 | NA | NA | NA |
| *Troglodytes troglodytes* | Northern Wren | 0.30 | 0.64 | 0.43 | 0.60 |
| *Turdus iliacus* | Redwing | 0.02 | 0.06 | NA | NA |
| *Turdus merula* | Eurasian Blackbird | -0.03 | 0.11 | 0.07 | 0.17 |
| *Turdus philomelos* | Song Thrush | 0.06 | 0.15 | 0.08 | 0.25 |
| *Turdus pilaris* | Fieldfare | 0.18 | 0.16 | NA | NA |
| *Turdus torquatus* | Ring Ouzel | 0.09 | NA | 0.07 | NA |
| *Turdus viscivorus* | Mistle Thrush | NA | 0.07 | 0.003 | 0.17 |
| *Vanellus vanellus* | Northern Lapwing | NA | 0.02 | NA | 0.05 |

### Appendix 3. Estimated mean spatial population synchrony in log abundance for each species by country. *NA* indicates that the species was not in the country associated with the column. Species names in Latin and English common names provided.

| **Species** | **Common Name** | **Norway** | **Sweden** | **Switzerland** | **United Kingdom** |
| --- | --- | --- | --- | --- | --- |
| *Acanthis flammea* | Common Redpoll | 0.50 | NA | NA | NA |
| *Aegithalos caudatus* | Long-tailed Tit | NA | NA | NA | 0.25 |
| *Alauda arvensis* | Eurasian Skylark | NA | NA | 0.13 | 0.38 |
| *Anas platyrhynchos* | Mallard | NA | 0.01 | NA | 0.17 |
| *Anthus pratensis* | Meadow Pipit | 0.48 | NA | NA | 0.01 |
| *Anthus spinoletta* | Water Pipit | NA | NA | 0.21 | NA |
| *Anthus trivialis* | Tree Pipit | 0.41 | 0.02 | 0.10 | NA |
| *Apus apus* | Common Swift | NA | -0.009 | -0.02 | 0.42 |
| *Ardea cinerea* | Grey Heron | NA | NA | NA | 0.30 |
| *Branta canadensis* | Canada Goose | NA | -0.01 | NA | NA |
| *Bucephala clangula* | Common Goldeneye | NA | -0.02 | NA | NA |
| *Buteo buteo* | Eurasian Buzzard | NA | NA | 0.14 | 0.51 |
| *Carduelis cannabina* | Common Linnet | NA | NA | NA | 0.27 |
| *Carduelis carduelis* | European Goldfinch | NA | NA | 0.28 | 0.76 |
| *Certhia brachydactyla* | Short-toed Treecreeper | NA | NA | 0.11 | NA |
| *Certhia familiaris* | Eurasian Treecreeper | NA | 0.27 | 0.49 | NA |
| *Chloris chloris* | European Greenfinch | 0.03 | 0.47 | 0.41 | 0.61 |
| *Chroicocephalus ridibundus* | Black-headed Gull | NA | -0.02 | NA | NA |
| *Columba livia* | Rock Dove | NA | NA | NA | 0.09 |
| *Columba oenas* | Stock Dove | NA | NA | NA | 0.04 |
| *Columba palumbus* | Common Woodpigeon | 0.22 | 0.04 | 0.61 | 0.55 |
| *Corvus corax* | Common Raven | 0.17 | 0.10 | NA | NA |
| *Corvus corone* | Carrion Crow | 0.16 | 0.08 | 0.35 | 0.21 |
| *Corvus frugilegus* | Rook | NA | NA | NA | 0.06 |
| *Corvus monedula* | Western Jackdaw | NA | 0.07 | NA | 0.50 |
| *Cuculus canorus* | Common Cuckoo | 0.18 | 0.02 | 0.04 | 0.42 |
| *Cyanistes caeruleus* | Eurasian Blue Tit | NA | 0.10 | 0.50 | 0.36 |
| *Delichon urbicum* | Common House Martin | NA | 0.02 | NA | 0.14 |
| *Dendrocopos major* | Great Spotted Woodpecker | NA | 0.22 | 0.47 | 0.79 |
| *Dryocopus martius* | Black Woodpecker | NA | 0.15 | NA | NA |
| *Emberiza citrinella* | Yellowhammer | 0.08 | 0.43 | NA | 0.16 |
| *Emberiza schoeniclus* | Common Reed Bunting | 0.29 | 0.09 | NA | NA |
| *Erithacus rubecula* | European Robin | 0.43 | 0.21 | 0.52 | 0.48 |
| *Falco tinnunculus* | Common Kestrel | NA | NA | NA | 0.46 |
| *Ficedula hypoleuca* | European Pied Flycatcher | 0.38 | 0.18 | NA | NA |
| *Fringilla coelebs* | Common Chaffinch | 0.35 | 0.10 | 0.36 | 0.16 |
| *Fringilla montifringilla* | Brambling | 0.25 | NA | NA | NA |
| *Gallinago gallinago* | Common Snipe | 0.26 | NA | NA | NA |
| *Gallinula chloropus* | Common Moorhen | NA | NA | NA | 0.18 |
| *Garrulus glandarius* | Eurasian Jay | NA | 0.03 | 0.23 | 0.22 |
| *Grus grus* | Common Crane | NA | 0.14 | NA | NA |
| *Hirundo rustica* | Barn Swallow | NA | 0.006 | 0.12 | 0.36 |
| *Lagopus lagopus* | Willow Ptarmigan | 0.37 | NA | NA | NA |
| *Lagopus muta* | Rock Ptarmigan | 0.38 | NA | NA | NA |
| *Larus argentatus* | European Herring Gull | NA | -0.03 | NA | 0.14 |
| *Larus canus* | Common Gull | NA | -0.02 | NA | NA |
| *Larus fuscus* | Lesser Black-backed Gull | NA | NA | NA | 0.26 |
| *Lophophanes cristatus* | Crested Tit | NA | 0.21 | 0.17 | NA |
| *Loxia curvirostra* | Red Crossbill | NA | 0.23 | NA | NA |
| *Lyrurus tetrix* | Black Grouse | 0.006 | 0.23 | NA | NA |
| *Motacilla alba* | White Wagtail | 0.16 | 0.12 | 0.11 | 0.25 |
| *Muscicapa striata* | Spotted Flycatcher | 0.21 | 0.02 | NA | NA |
| *Oenanthe oenanthe* | Northern Wheatear | 0.24 | NA | 0.21 | NA |
| *Parus major* | Great Tit | 0.41 | 0.03 | 0.36 | 0.72 |
| *Passer domesticus* | House Sparrow | NA | 0.01 | 0.18 | 0.06 |
| *Passer montanus* | Eurasian Tree Sparrow | NA | 0.03 | 0.07 | NA |
| *Periparus ater* | Coal Tit | NA | 0.05 | 0.24 | 0.23 |
| *Phasianus colchicus* | Common Pheasant | NA | NA | NA | 0.46 |
| *Phoenicurus ochruros* | Black Redstart | NA | NA | 0.50 | NA |
| *Phoenicurus phoenicurus* | Common Redstart | 0.40 | 0.05 | NA | NA |
| *Phylloscopus bonelli* | Western Bonelli's Warbler | NA | NA | 0.29 | NA |
| *Phylloscopus collybita* | Common Chiffchaff | 0.47 | NA | 0.53 | 0.66 |
| *Phylloscopus sibilatrix* | Wood Warbler | NA | 0.07 | NA | NA |
| *Phylloscopus trochilus* | Willow Warbler | 0.53 | 0.09 | NA | 0.33 |
| *Pica pica* | Eurasian Magpie | NA | 0.07 | 0.27 | 0.04 |
| *Picus viridis* | European Green Woodpecker | NA | NA | NA | 0.08 |
| *Pluvialis apricaria* | European Golden Plover | 0.03 | NA | NA | NA |
| *Poecile montanus* | Willow Tit | 0.07 | 0.18 | NA | NA |
| *Poecile palustris* | Marsh Tit | NA | NA | 0.10 | NA |
| *Prunella collaris* | Alpine Accentor | NA | NA | 0.02 | NA |
| *Prunella modularis* | Dunnock | 0.33 | 0.11 | 0.11 | 0.35 |
| *Pyrrhula pyrrhula* | Eurasian Bullfinch | NA | NA | 0.15 | 0.20 |
| *Regulus ignicapilla* | Common Firecrest | NA | NA | 0.46 | NA |
| *Regulus regulus* | Goldcrest | NA | 0.53 | 0.36 | 0.34 |
| *Saxicola rubetra* | Whinchat | NA | 0.03 | NA | NA |
| *Serinus serinus* | European Serin | NA | NA | 0.13 | NA |
| *Sitta europaea* | Eurasian Nuthatch | NA | 0.26 | 0.20 | NA |
| *Spinus spinus* | Eurasian Siskin | 0.39 | 0.35 | NA | NA |
| *Streptopelia decaocto* | Eurasian Collared Dove | NA | NA | NA | 0.14 |
| *Sturnus vulgaris* | Common Starling | NA | 0.03 | 0.17 | 0.47 |
| *Sylvia atricapilla* | Eurasian Blackcap | NA | 0.29 | 0.53 | 0.74 |
| *Sylvia borin* | Garden Warbler | NA | 0.14 | 0.16 | NA |
| *Sylvia communis* | Common Whitethroat | NA | 0.09 | NA | 0.55 |
| *Sylvia curruca* | Lesser Whitethroat | NA | 0.30 | NA | NA |
| *Tringa ochropus* | Green Sandpiper | NA | 0.13 | NA | NA |
| *Tringa totanus* | Common Redshank | 0.18 | NA | NA | NA |
| *Troglodytes troglodytes* | Northern Wren | 0.69 | 0.73 | 0.47 | 0.62 |
| *Turdus iliacus* | Redwing | 0.43 | 0.35 | NA | NA |
| *Turdus merula* | Eurasian Blackbird | 0.16 | 0.09 | 0.17 | 0.40 |
| *Turdus philomelos* | Song Thrush | 0.27 | 0.17 | 0.38 | 0.36 |
| *Turdus pilaris* | Fieldfare | 0.21 | 0.20 | NA | NA |
| *Turdus torquatus* | Ring Ouzel | 0.28 | NA | 0.18 | NA |
| *Turdus viscivorus* | Mistle Thrush | NA | 0.19 | 0.11 | 0.39 |
| *Vanellus vanellus* | Northern Lapwing | NA | 0.06 | NA | 0.26 |

### Appendix 4. Parameter estimates and standard errors (parentheses) for all chosen models across all distance intervals for log population growth rate (A) and log abundance (B). Max distance varied by country, for max distance values see Table 1.

| **A.    Log population growth rate** | **Top Model** | | | |
| --- | --- | --- | --- | --- |
| **Parameter** | **0 - 350km** | **0 - 500km** | **0 - 1000km** | **0 – Max Distance** |
| Norway, Long-distance migrant | 0.29 (0.03) | 0.29 (0.03) | 0.25 (0.03) | 0.18 (0.03) |
| Norway, Short-distance migrant | 0.34 (0.03) | 0.32 (0.03) | 0.29 (0.03) | 0.25 (0.03) |
| Norway, Resident | 0.32 (0.03) | 0.32 (0.04) | 0.29 (0.04) | 0.22 (0.03) |
| Sweden, Long-distance migrant | 0.27 (0.03) | 0.22 (0.03) | 0.21 (0.03) | 0.21 (0.03) |
| Sweden, Short-distance migrant | 0.33 (0.03) | 0.32 (0.03) | 0.32 (0.03) | 0.29 (0.03) |
| Sweden, Resident | 0.30 (0.03) | 0.32 (0.03) | 0.30 (0.03) | 0.25 (0.03) |
| Switzerland, Long-distance migrant | 0.26 (0.03) | 0.28 (0.04) | 0.28 (0.03) | 0.22 (0.03) |
| Switzerland, Short-distance migrant | 0.32 (0.03) | 0.30 (0.03) | 0.31 (0.03) | 0.29 (0.03) |
| Switzerland, Resident | 0.30 (0.03) | 0.31 (0.03) | 0.32 (0.03) | 0.26 (0.03) |
| United Kingdom, Long-distance migrant | 0.36 (0.03) | 0.39 (0.04) | 0.36 (0.04) | 0.27 (0.03) |
| United Kingdom, Short-distance migrant | 0.42 (0.03) | 0.39 (0.03) | 0.37 (0.03) | 0.34 (0.03) |
| United Kingdom, Resident | 0.40 (0.03) | 0.34 (0.03) | 0.32 (0.03) | 0.31 (0.03) |
| Log Generation Time | -0.11 (0.02) | -0.11 (0.02) | -0.11 (0.02) | -0.12 (0.02) |

| **B.    Log abundance** | **Top Model** | | | |
| --- | --- | --- | --- | --- |
| **Parameter** | **0 - 350km** | **0 - 500km** | **0 - 1000km** | **0 – Max Distance** |
| Norway, Long-distance migrant | 0.40 (0.04) | 0.39 (0.04) | 0.37 (0.04) | 0.37 (0.04) |
| Norway, Short-distance migrant | 0.49 (0.04) | 0.48 (0.04) | 0.46 (0.04) | 0.48 (0.04) |
| Norway, Resident | 0.43 (0.04) | 0.42 (0.04) | 0.41 (0.04) | 0.42 (0.05) |
| Sweden, Long-distance migrant | 0.29 (0.04) | 0.28 (0.04) | 0.27 (0.04) | 0.25 (0.04) |
| Sweden, Short-distance migrant | 0.38 (0.04) | 0.37 (0.04) | 0.36 (0.04) | 0.36 (0.05) |
| Sweden, Resident | 0.32 (0.04) | 0.31 (0.04) | 0.31 (0.04) | 0.30 (0.05) |
| Switzerland, Long-distance migrant | 0.35 (0.04) | 0.35 (0.04) | 0.35 (0.04) | 0.34 (0.04) |
| Switzerland, Short-distance migrant | 0.45 (0.04) | 0.45 (0.04) | 0.45 (0.04) | 0.45 (0.04) |
| Switzerland, Resident | 0.39 (0.04) | 0.39 (0.04) | 0.39 (0.04) | 0.39 (0.04) |
| United Kingdom, Long-distance migrant | 0.54 (0.04) | 0.50 (0.04) | 0.46 (0.04) | 0.46 (0.05) |
| United Kingdom, Short-distance migrant | 0.63 (0.04) | 0.59 (0.04) | 0.55 (0.04) | 0.57 (0.05) |
| United Kingdom, Resident | 0.57 (0.04) | 0.53 (0.04) | 0.50 (0.04) | 0.50 (0.04) |
| Log Generation Time | -0.12 (0.03) | -0.12 (0.03) | -0.12 (0.03) | -0.14 (0.03) |

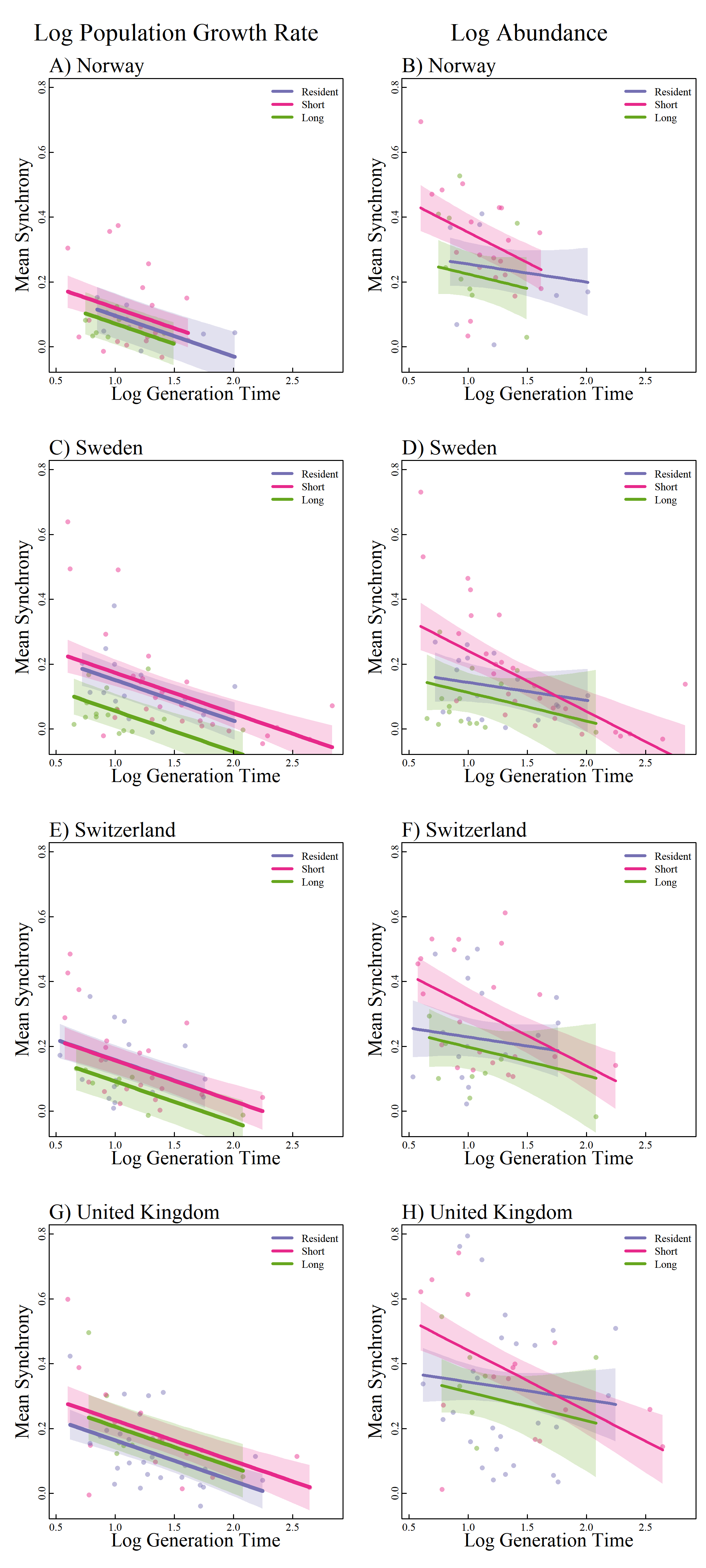

### Appendix 5. Interaction terms between migration tactic and country (log population growth rate) or migration tactic and generation time (log abundance) appeared in the top model. Mean synchrony is estimated from the log population growth rate top model (Country * Migration Tactic + Log Generation Time) and 0-max distance interval log abundance top model (Country + Migration Tactic * Log Generation Time). 95% confidence intervals are presented as shaded colors.
